# Urea excreted by tropical frogs is explained by their body mass and the mass of insects consumed

**DOI:** 10.1101/2023.10.09.561504

**Authors:** Akash M Dev, Gayathri Sreedharan, Yashwant Singh Panwar, Karthikeyan Vasudevan

## Abstract

Frog insect interactions are prominent and they result in services that are crucial in the ecosystem. The primary nitrogenous waste that frogs produce is urea. However, the role of biotic variables on urea excretion, has not been fully investigated. Frogs comprising of six *Polypedates maculatus* and seven *Minervarya agricola* were maintained in controlled condition after tagging them with passive integrated transponders. Thirteen experiments were performed to test whether, frog species or frogs’ body mass or insect body mass, or a combination of these explained the amount of urea excreted by the frogs. We found that body mass of the frogs and the amount of insect biomass consumed explained the amount of urea excreted in the two species. In the light of these findings, we discuss the role of frogs in converting insect biomass into urea in the ecosystem.

## INTRODUCTION

Arthropod biomass dominates the terrestrial biome as they account for 50% (1 Gt C) of the total animal biomass (Bar-On et al., 2018). Amphibian biomass depends on arthropod biomass, and it is one by tenth of the insect biomass (Bar-On et al., 2018). Frogs are amphibians and they constitute a diverse group that occur in a majority of the ecoregions of the world. Adult frogs are insectivores and they primarily feed on the arthropod biomass (Wells, 2010). Through this, they also regulate the abundance of terrestrial and aquatic arthropods (Wells, 2010) and insect pests (Springborn et al., 2022). In addition to this, a lesser known role is the excretion of urea by frogs (Teng et al., 2016). This could be an important source of nitrogen for plants in both terrestrial and freshwater ecosystems.

Trophic interactions of frogs and insects in the ecosystem might be important for ecosystem functioning. Therefore, it has been hypothesized that these services might have been severely impacted by enigmatic declines in frog populations (Gibbons et al., 2000). The declines caused by over-exploitation, habitat loss (Carey et al., 1999) and disease (Scheele et al., 2014) had devastating impact frog population worldwide. However, a thorough evaluation of the impacts of the amphibian declines on ecosystems have not been made. Such evaluation might strengthen the prospects of restoring frog populations and improving the ecosystem function.

Urea is excreted as the principal nitrogenous waste in frogs (Wright & Wright, 1996). However, the factors that influence excretion of urea have not been studied. Our aim was to estimate urea excreted by frogs when a known quantity of insects was fed under controlled abiotic conditions. Based on the data collected we tested whether body weight of frogs, weight of insect, the species of frog or a combination of these factors influence urea excreted?

## MATERIALS AND METHODS

### Experimental Design

Species of frogs: *Polypedates maculatus* and *Minervarya agricola* were maintained in an amphibian facility at Laboratory for Conservation of Endangered Species, Hyderabad, India. The frogs were captured from the campus premises, maintained in terraria for the experiments. Permission from Institutional Animal Ethics Committee was obtained for the experiments (Project no: IAEC 82/22). The design of the terraria were as per the prescribed guidelines (Vasudevan et al., 2022). For the experiment, 13 frogs (six *P. maculatus* and seven *M. agricola*) were maintained in captivity after tagging them with Passive Integrated transponders (PIT) tags. We performed thirteen experimental replicates on the two species of frogs. All experimental replicates were identical, and it started with: housing individual frogs in the terraria; fasting them for four days prior to the experiment (see Figure 1). They were fed with live orthopteran insects that were routinely caught using an insect net from the campus. The facility maintained was main temperature of 26°C, with humidity levels consistently set at 60%. Additionally, a 10-hour dark period was implemented, complemented by the provision of cold lighting. We assumed that fasting period made them sufficiently hungry and prepared them to feed. The two species differed in size and habitat use. We assumed that there was no effect of body weight on the fasting levels and digestive physiology. We could not identify sex of the frog as they did not show sexual dimorphism. We assumed that age and sex of the frogs would not influence digestive physiology and excretion of urea in the frogs. Since, frogs are generalist insectivores, we assumed that the two species did not show any preference for prey items and that the insects fed upon were equally digestible by them. On the fifth day, each experimental frog was screened for PIT tags, IDs recorded and transferred into separate crystallisation dishes (Borosil^®^ 190 x 100 mm). Snout to vent length (SVL) was measured using vernier callipers and body mass using Pesola^™^ scale with accuracy of 0.05g. Into each container 100ml of tap water kept open for 24 hours was poured and closed with a perforated lid. Two wild caught, live orthopteran insects were weighed and provisioned to a frog on the first day of the experiment. We assumed that body weight of the orthopteran insect was an accurate measure of the nutritive content of the insect. At the end of the experiment, the enclosures were checked for live or dead insects. Two experimental controls were used: 1) container with 100ml of standing water and two grasshoppers, without frog; 2) container with 100ml of standing water without frog. The experimental setup was not disturbed for 48 hours. After this period, water samples were collected from each container avoiding any cross-contamination. Frogs were transferred back into the terraria where they were sourced from. Water samples were taken to the lab for quantification of urea, using diacetyl monoxime method (DAM).

**Fig 1:**
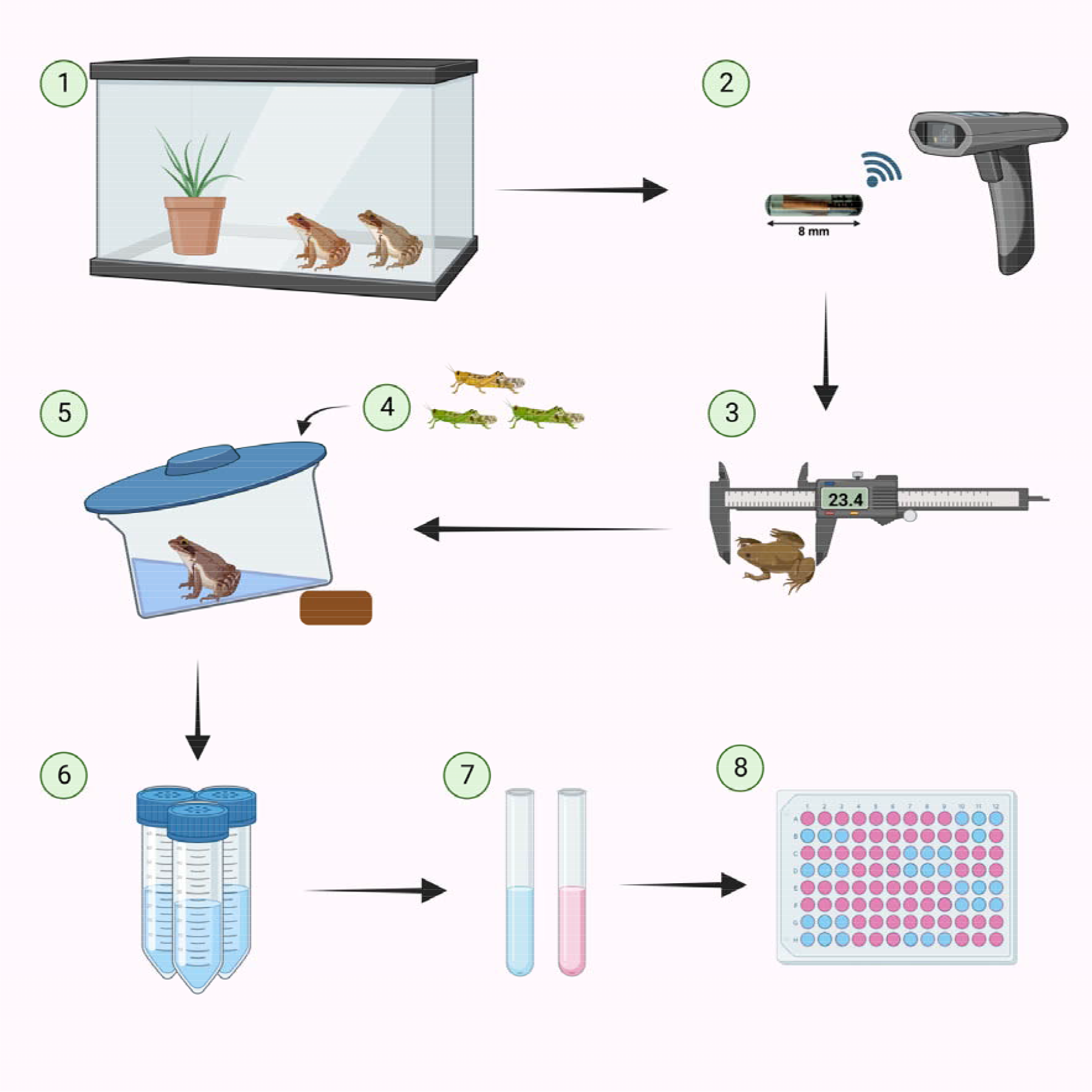
Steps involved in the experimental set-up: Frog species were captured and maintained in a terrarium at our facility to fast for 4 days (1). Each of them was given a separate passive integrated transponders tag (2) and was measured for snout vent length (3) and these were recorded before each experimental episode. After transferring the frogs to their individual containers (4), they were fed with grasshoppers on the day of the experiment (5). After 48 hrs, the water from the containers, were sampled for (5) urea estimation using DAM (7) method, in an ELISA (8) reader (7,8).

### DAM Method

We followed the protocol outlined in Bugbee (2021) and some modifications were made to it. (a) Acid reagent was prepared by dissolving 324mg ferric chloride in 10ml of 56% orthophosphoric acid. Then dilute ferric chloride and orthophosphoric acid mixture to 100ml of 20% sulphuric acid. (b) 0.62g of diacetyl monoxime (SIGMA^™^) was added and diluted to 100ml with distilled water in volumetric flasks. (c) 41mg thiosemicarbazide (NICE™) was added and diluted to 50ml with distilled water in volumetric flask. We prepared known concentrations of urea (RANKEM^™^) of 2x dilution ranging from 200 to 3.12 *μ*g/ml for standard curve. To all 20ml glass test tubes for sample, standards, and blank, 3ml of acid reagent, 1ml of Diacetyl monoxime and 1ml of thiosemicarbazide were added. 200 *μ*l of sample collected from each frog was added into 30ml test tubes and made up to 6 ml using double distilled water. Content of the test tube was vigorously mixed using Vortex-Genie 2 (Scientific Industries, Inc.^™^). Ensure the test tubes are labelled and kept in boiling water (Julabo^™^water bath) bath for 15 minutes and remove tubes from water bath and allowed it to cool for 10 minutes. From the tubes 300*μ*l of all samples were transferred to ELISA plate in triplicates and the absorbance was read at 520nm using ELISA plate reader (ThermoFisher^™^). Optical Density (OD) values were obtained in triplicates for each sample, an average OD value was taken for each sample. The value of blank was subtracted from the OD value of the standard as well as each sample and used for further calculation. Using standard curve, urea concentration of samples was determined.

### Data analyses

The data was subjected to Shapiro Wilk’s test for homogeneity variance. After checking for normality of distribution, generalized linear mixed modelling (GLMM) was used to test the hypotheses on the factors that influence urea excretion. In twelve out of thirteen experiments both insects added were consumed by the frog, and in one experiment an insect was found dead. Therefore, we assumed that difference in concentration of urea was not influenced by mortality of the insect during the experiment. Body weight of the experimental frogs, weight of the insect fed and species of frog were considered as random variable in the models. Since, the experiment was replicated 13 times, the ID of the experimental frog was considered as fixed effect in the models. Four models were constructed changing the random variables, and models with the lowest AIC values was considered, competing models that had AIC values within a difference of two were taken for model averaging was done (Beier et al., 2001). The model weights were considered as the variation explained. Model with highest weight was considered as the best fit model for the hypothesis. All statistical analyses including *lme4, lmerTest* and *MuMIn* analysis were performed using R studio (version 2022.02.1+461).

## RESULTS

The experiment lasted 91 days, and 7 *P. maculatus* and 6 *M. agricola* were recruited in the experiment. 4 of *P. maculatus* and 4 of *M. agricola* survived the entire duration of the experiment. Experimental frogs belonging to *P. maculatus* and *M. agricola* weighed 6g (SE = 0.17), and 3.87g (SE ± 0.26), respectively. They were significantly different in their body mass (t = 2.344, df = 11, P = 0.03). Mean weight of insects provisioned during the experiment for *P. maculatus* and *M. agricola* were 0.185g (SE ± 0.17), and 0.164g (SE ± 0.01), respectively. The mean change in body weight for *P. maculatus* and *M. agricola* were -0.97g (SE ± 0.47, t = -2.04, df = 6, P = 0.08) and 0.55g (SE ± 0.51, t = 1.06, df = 5, P = 0.33) per day, respectively. The concentration of urea excreted was significantly different from control (t =14.05, P = 0.02) The mean concentration of urea excreted per gram body mass of the frogs: *P. maculatus* and *M. agricola* were 24 *μ* g (SE ± 2) and 32 *μ* g (SE ± 2.85), respectively.

Out of the four mixed models, first three models had similar Akaike information criterion (AIC) values (Table 1). In them models frog weight was a random variable. In the model with the highest AIC value insect weight was a random variable. The first model was the global model, which had highest AIC value insect weight, frog weight and species as a random variable. After model averaging, weight of the insect and weight of the frog explained 47% of the variation in the amount of urea excreted by the frog in the experiments (Table 2).

**Table 1:** Description of the models with statistical parameters to evaluate them and explain urea excreted by frogs in the experiment.

| Model | Description | AIC | Deviance | Degree of freedom |
| --- | --- | --- | --- | --- |
| 1 | urea concentration ~ grasshopper weight + frog weight + species+(1\| tag id) | 978.9 | 966.92 | 6 |
| 2 | urea concentration ~ grasshopper weight+ frog weight + (1 \| tag id) | 978.3 | 968.33 | 5 |
| 3 | urea concentration ~ frog weight + (1 \|tag id) | 979.9 | 971.91 | 4 |
| 4 | urea concentration ~ grasshopper weight + (1 \| tag id) | 986.6 | 989.61 | 4 |

Abbreviation: AIC, The Akaike information criterion (AIC)

**Table 2:** Model averaging results on top three models that explained urea excreted by frogs in the experiment.

| Model rank | Description | Weight |
| --- | --- | --- |
| 1 | urea concentration~ grasshopper weight +frog weight + (1 \| tag id) | 0.47 |
| 2 | urea concentration~ grasshopper weight +frog weight + species + (1 \| tag id) | 0.29 |
| 3 | urea concentration ~ frog weight + (1 \| tag id) | 0.24 |

## DISCUSSION

Both *M. agricola* and *P. maculatus* are abundant in cultivated land and human modified areas. The former is a semi-aquatic and the latter is an arboreal frog. In natural ecosystems, insects are abundant and frogs feed *ad libitum*. Therefore, the estimate of urea excreted could be much higher than that estimated through this experiment. Physiological process of excretion in frogs has been extensively studied (Balinsky & Baldwin, 1961). These studies used invasive methods to decipher the urea cycle in frogs (Balinsky et al., 1976). The study of enzymes involved in the urea cycle during metamorphosis has revealed from the transition from ammonotelism to ureotelism excretion (Brown et al., 1959). Enhanced activity of enzymes such as, carbomyl phosphate synthase I, ornithine transcarbamylase and arginase have been documented (Helbing et al., 1992). The activity of these enzymes might influence urea excretion in frogs.

Frogs transfer nitrogen from the insect biomass in the form of urea, so that could be readily taken up by plants. The biomass of frogs and insects are therefore, important for transfer of nitrogen above ground to the soil. Understanding physiological factors influencing urea excretion is crucial to investigate the process of urea formation and its transfer to the environment. This study specifically quantifies the amount of urea excreted by frogs by insectivory, so that it might stimulate further investigation.

The microhabitats occupied by frogs might allow slow seeding and dispersion of urea in ways that might be important for terrestrial plants, including agricultural crops. Arboreal frogs are known to seed nutrients to epiphytic plants (Romero et al., 2010). Depending on species of frogs, their habitat and season, the prey would vary (Hirai & Matsui, 1999), and it might have an effect on the nutrient cycling in the ecosystem. The role of larval anurans (tadpoles) in nutrient cycling in the aquatic ecosystems is widely recognised (Seale & Beckvar, 1980, Connelly et al., 2008, Connelly et al., 2008, Whiles et al., 2013). Such role for adult frogs has not been recognised (Valencia-Aguilar et al., 2013; Hocking & Babbitt, 2014). Adult anurans occupy nearly the entire terrestrial biome, and they are usually terrestrial or semi-terrestrial. They are also voracious insectivores, and are agents of pest biological control (Abdulali, 1985; Seshadri et al., 2020). In the tropics, where the abundance of anurans is high, they might play an important role in mediating nutrient cycling in the forest floor (Beard et al., 2002). Nutrient cycling in the soil has wide ramifications for the ecosystem, as nitrogen in the soil inhibits enzymes responsible for recalcitrant carbon degradation and promotes carbon storage (Macdonald et al., 2018). As impoverishment of frogs (Luedtke et al., 2023) and insects (Wagner, 2020) continue globally, it might unleash a nutrient cascade on terrestrial ecosystems that has not yet been recognised. The enigmatic declines of amphibians (Stuart et al., 2004) for over one third of a century (Luedtke et al., 2023) might have already disrupted nutrient cycling in terrestrial ecosystems.

## Acknowledgement

The CSIR-CCMB supported this work providing funds and instrumentation facilities. All the faculty members from School of bioscience Mar Athanasios College for Advanced Studies, Tiruvalla supported AM for working with CCMB. Harika Segu, Vinod Kumar, K. Rajyalakshmi, provided assistance during the experiments. M Jagadeesh Yadav assisted in maintaining the terrarium. We also thank Ravi Kumar Singh and Bharath Subramanyam for their support throughout this work.

## References

Abdulali, H. (1985). On the export of frog legs from India. Journal of the Bombay Natural History Society. Bombay, 82(2), 347–375.

Balinsky, J. B., & Baldwin, E. (1961). The Mode of Excretion of Ammonia and Urea in Xenopus Laevis. Journal of Experimental Biology, 38(4), 695–705. 10.1242/jeb.38.4.695

Balinsky, J. B., Chemaly, S. M., Currin, A. E., Lee, A. R., Thompson, R. L., & Van der Westhuizen, D. R. (1976). A comparative study of enzymes of urea and uric acid metabolism in different species of Amphibia, and the adaptation to the environment of the tree frog Chiromantis xerampelina Peters. Comparative Biochemistry and Physiology -- Part B: Biochemistry And, 54(4), 549–555. 10.1016/0305-0491(76)90139-5

Bar-On, M.Y., Phillips, R., Milo, R. (2018). The biomass distribution on Earth. PNAS, 115 (25) 6506–6511. 10.1073/pnas.1711842115

Beard, K. H., Vogt, K. A., & Kulmatiski, A. (2002). Top-down effects of a terrestrial frog on forest nutrient dynamics. Oecologia, 133(4), 583–593. 10.1007/s00442-002-1071-9

Beier, P., Burnham, K. P., & Anderson, D. R. (2001). Model Selection and Inference: A Practical Information-Theoretic Approach. In The Journal of Wildlife Management (Vol. 65, Issue 3). 10.2307/3803117

Brown, G. W., Brown, W. R., & Cohen, P. P. (1959). Comparative Biochemistry of Urea Synthesis. Journal of Biological Chemistry, 234(7), 1775–1780. 10.1016/s0021-9258(18)69924-7

Carey, C., Cohen, N., & Rollins-Smith, L. (1999). Amphibian declines: An immunological perspective. Developmental and Comparative Immunology, 23(6), 459–472. 10.1016/S0145-305X(99)00028-2

Connelly, S., Pringle, C. M., Bixby, R. J., Brenes, R., Whiles, M. R., Lips, K. R., Kilham, S., & Huryn, A. D. (2008). Changes in stream primary producer communities resulting from large-scale catastrophic amphibian declines: Can small-scale experiments predict effects of tadpole loss?Ecosystems, 11(8), 1262–1276. 10.1007/s10021-008-9191-7

Gibbons, J. W., Scott, D. E., Ryan, T. J., Buhlmann, K. A., Tuberville, T. D., Metts, B. S., Greene, J. L., Mills, T., Leiden, Y., Poppy, S., & Winne, C. T. (2000). The Global Decline of Reptiles, Déjà Vu Amphibians: Reptile species are declining on a global scale. Six significant threats to reptile populations are habitat loss and degradation, introduced invasive species, environmental pollution, disease, unsustaina. BioScience, 50(8), 653–666. 10.1641/0006-3568(2000)050[0653:TGDORD]2.0.CO%0A http://0.0.0.2

Helbing, C., Gergely, G., & Atkinson, B. G. (1992). Sequential up[regulation of thyroid hormone β receptor, ornithine transcarbamylase, and carbamyl phosphate synthetase mRNAs in the liver of Rana catesbeiana tadpoles during spontaneous and thyroid hormone[induced metamorphosis. Developmental Genetics, 13(4), 289–301. 10.1002/dvg.1020130406

Hirai, T., & Matsui, M. (1999). Feeding habits of the pond frog, Rana nigromaculata, inhabiting rice fields in Kyoto, Japan. Copeia, 1999(4), 940–947. 10.2307/1447969

Hocking, D. J., & Babbitt, K. J. (2014). University of New Hampshire Scholars ‘Repository Amphibian Contributions to Ecosystem Services. 9, 1–17.

Luedtke, J. A., Chanson, J., Neam, K., Hobin, L., Maciel, A. O., Catenazzi, A., Borzée, A., Hamidy, A., Aowphol, A., Jean, A., Sosa-Bartuano, Á., Fong G. A., de Silva, A., Fouquet, A., Angulo, A., Kidov, A. A., Muñoz Saravia, A., Diesmos, A. C., Tominaga, A., … Stuart, S. N. (2023). Ongoing declines for the world’s amphibians in the face of emerging threats. Nature. 10.1038/s41586-023-06578-4

Macdonald, C. A., Delgado-Baquerizo, M., Reay, D. S., Hicks, L. C., & Singh, B. K. (2018). Soil nutrients and soil carbon storage: Modulators and mechanisms. In Soil Carbon Storage: Modulators, Mechanisms and Modeling. Elsevier Inc. 10.1016/B978-0-12-812766-7.00006-8

Romero, G. Q., Nomura, F., Gonçalves, A. Z., Dias, N. Y. N., Mercier, H., Conforto, E. de C., & Rossa-Feres, D. de C. (2010). Nitrogen fluxes from treefrogs to tank epiphytic bromeliads: An isotopic and physiological approach. Oecologia, 162(4), 941–949. 10.1007/s00442-009-1533-4

Scheele, B. C., Hunter, D. A., Grogan, L. F., Berger, L., Kolby, J. E., Mcfadden, M. S., Marantelli, G., Skerratt, L. F., & Driscoll, D. A. (2014). Interventions for Reducing Extinction Risk in Chytridiomycosis-Threatened Amphibians. Conservation Biology, 28(5), 1195–1205. 10.1111/cobi.12322

Seale, D. B., & Beckvar, N. (1980). The Comparative Ability of Anuran Larvae (Genera: Hyla, Bufo and Rana) to Ingest Suspended Blue-Green Algae. Copeia, 1980(3), 495–503. 10.2307/1444527

Seshadri, K. S., Allwin, J., Seena, N. K., & Ganesh, T. (2020). Anuran assemblage and its trophic relations in rice-paddy fields of South India. Journal of Natural History, 54(41– 42), 2745–2762. 10.1080/00222933.2020.1867772

Springborn, M. R., Weill, J. A., Lips, K. R., Ibáñez, R., & Ghosh, A. (2022). Amphibian collapses increased malaria incidence in Central America. Environmental Research Letters, 17(10). 10.1088/1748-9326/ac8e1d

Stuart, S. N., Chanson, J. S., Cox, N. A., Young, B. E., Rodrigues, A. S. L., Fischman, D. L., & Waller, R. W. (2004). Status and trends of amphibian. Science, 306(5702), 1783–1786.

Teng, Q., Hu, X. F., Luo, F., Cheng, C., Ge, X., Yang, M., & Liu, L. (2016). Influences of introducing frogs in the paddy fields on soil properties and rice growth. Journal of Soils and Sediments, 16(1), 51–61. 10.1007/s11368-015-1183-6

Valencia-Aguilar, A., Cortés-Gómez, A. M., & Ruiz-Agudelo, C. A. (2013). Ecosystem services provided by amphibians and reptiles in Neotropical ecosystems. International Journal of Biodiversity Science, Ecosystem Services and Management, 9(3), 257–272. 10.1080/21513732.2013.821168

Vasudevan, K., Barkha, S., Venu, G., Gowri, M., Lisa, G., & Kishor, G. B. (2022). Ex-situ management of amphibians in Indian zoos, Central Zoo Authority, New Delhi.

Wagner, D. L. (2020). Insect declines in the anthropocene. Annual Review of Entomology, 65, 457–480. 10.1146/annurev-ento-011019-025151

Wells, K. D. (2010). The ecology and behavior of amphibians. In The Ecology and Behavior of Amphibians. University of Chicago press.

Whiles, M. R., Hall, R. O., Dodds, W. K., Verburg, P., Huryn, A. D., Pringle, C. M., Lips, K. R., Kilham, S. S., Colón-Gaud, C., Rugenski, A. T., Peterson, S., & Connelly, S. (2013). Disease-Driven Amphibian Declines Alter Ecosystem Processes in a Tropical Stream. Ecosystems, 16(1), 146–157. 10.1007/s10021-012-9602-7

Wright, P. M., & Wright, P. A. (1996). Nitrogen metabolism and excretion in bullfrog (Rana catesbeiana) tadpoles and adults exposed to elevated environmental ammonia levels. Physiological Zoology, 69(5), 1057–1078. 10.1086/physzool.69.5.30164246

